# PEELING WALLS1 encodes a GT106 protein required for seed surface integrity

**DOI:** 10.64898/2026.09.18.752628

**Authors:** Alan D. Gomez Vargas, Talia Jacobson, Cătălin Voiniciuc

## Abstract

Rhamnogalacturonan I (RG-I) is a major pectin domain and an important wall component for cell-cell adhesion. Arabidopsis seed mucilage serves as a powerful model to identify and characterize pectin-related enzymes, since this gelatinous capsule is composed predominantly of RG-I. Although multiple glycosyltransferase (GT) families participate in pectin biosynthesis, only a limited number of their members have been functionally characterized *in vivo* or biochemically characterized *in vitro*. In this study, we characterized the biological functions of PEELING WALLS1 (PEEL1), a Golgi-localized GT106 protein that is related to known RG-I rhamnosyltransferases (RRTs). Knocking out *PEEL1*, but not its close paralog *PEEL1-LIKE* (*PEEL1L*), led to patchy mucilage release from seeds upon hydration. However, RG-I content was not compromised in mucilage extracted from *peel1* or *peel1 peel1L* mutant seeds. Treatment of seeds with a cation chelator restored mucilage expansion but failed to rescue the underlying defect in epidermal cell adhesion. Histological experiments and transgene complementation demonstrated that optimal expression of *PEEL1* is required for the adhesion of primary walls to the seed surface. Collectively, these findings identify *PEEL1* as a previously unrecognized contributor to primary cell wall adhesion and suggest that different RRT-related proteins contribute to unique pectic structures or localizations *in vivo*, rather than total mucilage RG-I abundance.

## Introduction

Seed coat cells differentiate from the ovule integument to form a protective interface between the embryo and the external environment. During this process, the polar deposition of copious amounts of pectin and some (hemi)cellulosic fibers in apoplastic pockets leads to the formation of gelatinous seed mucilage (Haughn and Chaudhury, 2005; Voiniciuc et al., 2015c). Upon hydration of dry seeds, mucilage rapidly expands and ruptures the primary cell wall to release a hydrogel that encapsulates the seed surface. Mutants with histological and/or biochemical defects in seed mucilage have enabled the characterization of Carbohydrate-Active enzymes (CAZymes) affecting polysaccharides structure and function (Voiniciuc et al., 2015a; Voiniciuc et al., 2015b; Voiniciuc et al., 2018; Saez-Aguayo et al., 2025).

Pectins are a diverse group of acidic polysaccharides that are enriched in primary cell walls and the middle lamella, where they contribute to cell-cell adhesion, porosity, and ion exchange capacity (Cosgrove, 2024). A large repertoire of glycosyltransferases (GTs) is required to synthesize the main domains of pectin: homogalacturonan (HG), rhamnogalacturonan I (RG-I), and RG-II. HG is a linear polymer containing only α-1,4-D-galacturonic acid (GalA), whose solubility and enzyme accessibility can be modified by methyl or acetyl ester groups. The RG-I backbone has alternating GalA and α-1,2-L-rhamnose (Rha) units, which can be decorated with long side chains such as arabinans, galactans or arabinogalactans. In contrast, RG-II features a HG backbone with complex branches composed of a dozen sugars, including rare residues.

In *Arabidopsis thaliana* (hereafter Arabidopsis), RG-I accounts for ∼90% of seed mucilage, with smaller amounts of HG and hemicelluloses (Voiniciuc et al., 2015c). This system has enabled the identification of key enzymes for pectin synthesis, such as *MUCILAGE-RELATED70* (*MUCI70*) and GALACTURONOSYLTRANFERASE11 (GAUT11). Mutations in *GAUT11*, encoding a GT8 enzyme that elongates HG *in vitro*, partly impaired mucilage release (Voiniciuc et al., 2018). In contrast, impaired RG-I accumulation in *muci70* mutants had more severe effects on mucilage release, which was nearly eliminated for *gaut11 muci70* double mutant seeds (Voiniciuc et al., 2018). Subsequently, recombinant MUCI70 was shown to have RG-I galacturonosyltransferase (RGGAT) activity and became the founding member of a novel GT116 family (Amos et al., 2022).

Along with MUCI70/RGGAT, mucilage synthesis requires RG-I rhamnosyltransferases (RRTs) from the GT106 family. Of the 34 GT106 genes in Arabidopsis, four RRT isoforms (RRT1-4) were initially shown to have redundant catalytic activity, with six additional enzymes forming two neighboring clades (**Fig. 1A**). Mutations in *RRT1* partly reduced mucilage Rha and GalA content and mucilage capsule size (Takenaka et al., 2018). Recently, FRIABLE1 (FRB1) protein, a protein related to RRT1-4, was shown to be an active RRT *in vitro* (Robichaux et al., 2026). RG-I oligosaccharides could be elongated when pairing RRT1 or FRB1 with MUCI70/RGGAT, suggesting cooperative or redundant roles in certain GT106 clades. More than a decade before these biochemical assays, *frb1* seedlings were shown to have impaired cell-cell adhesion, without altering overall pectin content (Neumetzler et al., 2012). In addition to the pectin-related roles, the GT106 family includes at least two groups of enzymes important for hemicellulose backbone elongation. MANNAN SYNTHESIS-RELATED (MSR) proteins enhance mannan synthases (Wang et al., 2013; Voiniciuc et al., 2019), while FRAGILE FIBER9 (FRA9) affects xylan content in the stem (Zhong et al., 2023).

**Figure 1.**
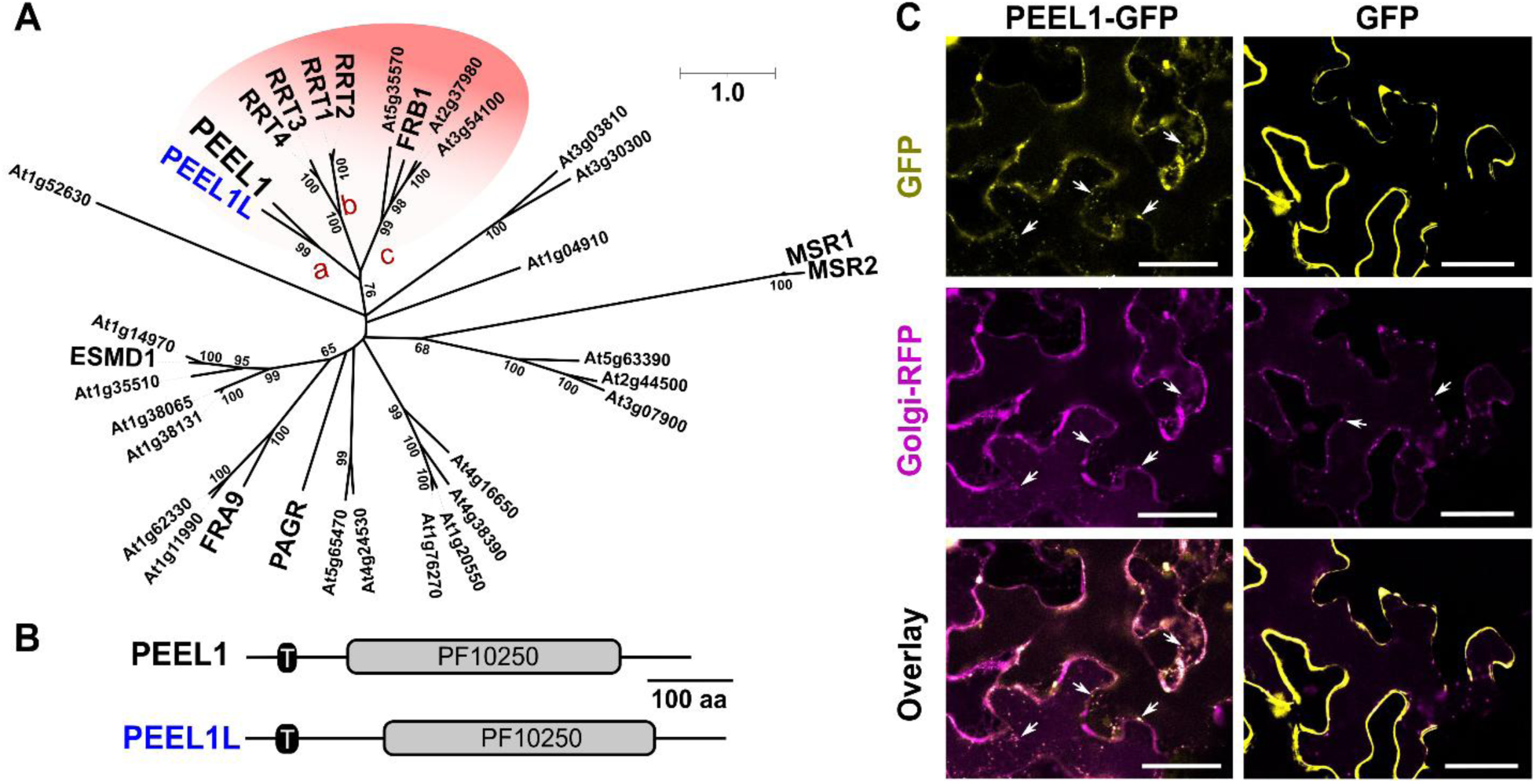
PEEL1 and PEEL1L are related to RRT proteins. **(A)** Phylogenetic tree of the 34 Arabidopsis GT106 members. RRT cluster includes PEEL1/1L (a), RRT1-4 (b), and third subclade containing FRB1 (c). Numbers show bootstrap values > 50. **(B)** Schematic of PEEL1/1L proteins. T, transmembrane domain. **(C)** PEEL1-GFP co-localizes with the Golgi marker GmMan1-mCherry, unlike the untagged GFP control, in *N. benthamiana* leaf cells. White arrows show punctae consistent with Golgi bodies. Scale bar = 50 µm. **Alt text:** Figure 1. Phylogenetic relationships, protein domain organization, and subcellular localization of PEEL protein. (A) Phylogenetic tree illustrating the evolutionary relationship between PEEL1, PEEL1L, and RRT proteins in Arabidopsis thaliana. (B) Schematic representation of PEEL1 and PEEL1L proteins, highlighting a transmembrane region followed by a catalytic domain. (C) Confocal microscopy images showing the co-localization of PEEL1-GFP with the Golgi apparatus-marker and a cytosolic GFP as control.

In this report, we functionally characterized an *RRT*-like gene named *PEELING WALLS1* (*PEEL1*). We demonstrate that Golgi-localized PEEL1 modulates pectin gelling properties and seed surface integrity,. A *peel1* transcriptional knockout mutant displayed patchy mucilage release without altering total mucilage composition. Histological experiments revealed underlying defects in the attachment of primary cell walls to the seed surface, which requires optimal expression of *PEEL1*, but not its paralog *PEEL1-LIKE* (*PEEL1L*).

## Materials and methods

### Plant material

Arabidopsis Col-0 wild-type (WT) and T-DNA insertional lines were cultivated in growth chambers set to have constant light, humidity and temperature, as previously reported (Jacobson et al., 2026). Mutations in At5g64600 (*peel1*) and At1g22460 (*peel1L*) were obtained from the GABI-DUPLO collection (Pair ID 2655) via the NASC stock center (https://arabidopsis.info/, Bolle et al., 2013). Mutants were genotyped using specific primers **(Supplementary Table 1)** according to the GABI-Kat design tool (https://www.gabi-kat.de/db/primerdesign.php). For transient expression, *Nicotiana benthamiana* plants were grown in a growth chamber with a 16:8h photoperiod.

### Bioinformatic analyses

Arabidopsis GT106 proteins phylogenetic relationships were inferred using the IQ-TREE web server (http://iqtree.cibiv.univie.ac.at/). GT106 amino acid sequences were retrieved from TAIR (https://www.arabidopsis.org/) and the phylogenetic tree was constructed with the best substitution model (WAG+F+I), including all sites. Reliability was evaluated with the bootstrap method with 500 replicates. Protein domains were identified using the InterPro database (https://www.ebi.ac.uk/interpro/). AlphaFold3 (https://alphafoldserver.com/) protein models were compared in ChimeraX (https://www.cgl.ucsf.edu/chimerax/).

### RNA isolation and quantitative real-time PCR

Total RNA was isolated from four mature green siliques collected from the main stem of 5-week-old Arabidopsis plants using the Monarch Spin RNA Isolation Kit (New England Biolabs, MA, USA), following the manufacturer’s protocol. For each sample, 200 ng of total RNA was used for the RevertAid First Strand cDNA Synthesis Kit (Thermo Fisher Scientific, MA, USA) and random hexamer primers. Quantitative real-time PCR (qPCR) was performed on a CFX Duet Real-Time System using cDNA, SsoAdvanced Universal SYBR Green Supermix (Bio-Rad, CA, USA) and gene-specific primer pairs **(Supplementary Table 1)**. Transcript levels were calculated using the 2^-ΔΔCt^ method, relative to the geometric mean of *AtUBQ5* and *AtGAPC1* reference genes.

### Transgene complementation experiments

Plant expression vectors were assembled using modular cloning (MoClo) parts available from Addgene (Weber et al., 2011; Engler et al., 2014), and a MoClo-optimized *PEEL1* sequence synthesized for compatibility with C-terminal fusions **(Supplementary Table 2)**. MoClo Level 1 constructs to express PEEL1 fused to green fluorescent protein (GFP) were assembled with either the constitutive cauliflower mosaic virus 35S (CaMV35S) or the stronger *TESTA-ABUNDANT2* (*TBA2*) promoter (McGee et al., 2019). Constructs without GFP were cloned by introducing a STOP codon at the end of *PEEL1* **(Supplementary Table 1),** and by amplifying *PEEL1L* with its native STOP codon from Arabidopsis cDNA. Level M constructs contained the L1 parts plus the FAST-RFP fluorescence marker. After whole-plasmid sequencing, Level M vectors were introduced into *Agrobacterium tumefaciens* GV3101 and then transformed into Arabidopsis WT or *peel1* plants via floral dip.

### Histological analysis of seed mucilage

Mucilage was stained with 0.01% Ruthenium Red (RR) after hydrating mature seeds in water 10 min at 150 rpm using a microplate shaker (Thermo Scientific, Model 88882005). Seeds were then rinsed with water to remove residual dye. Alternatively, dry seeds were pre-treated with 50 mM EDTA (ethylenediaminetetraacetic acid, pH 9.5) for 60 min at 125 rpm. Seeds were then rinsed with water and stained with 0.01% RR. Images were acquired using a Axiocam 208 color camera on stereomicroscope (Zeiss).

Unstained primary cell walls and birefringence were assessed after gently hydrating seeds in water, transferring them to well slides, analyzing them with brightfield or an analyzer/polarizer module on a Axiovert A1 inverted microscope with Axiocam 202 mono camera (Zeiss). The same microscope and camera setup were used for epifluorescence after seed staining with 0.01% propidium iodide (PI).

### Confocal microscopy

For protein localization, *35S:PEEL1-GFP* was co-expressed with the known Golgi marker *Glycine max* α-1,2-mannosidase1 (GmMan1) fused to mCherry (G-rb/CD3-968; Nelson et al., 2007). *A. tumefaciens* cultures carrying each construct were resuspended in infiltration media (10 mM MgCl_2_, 10 mM MES pH 5.6, and 100 mM acetosyringone). Desired constructs, including the P19 silencing suppressor, were mixed to a final OD_600_ of 0.5 and injected into abaxial *N. benthamiana* leaf cells. Images were acquired at 3 days post-infiltration.

Confocal microscopy was performed on a Leica Stellaris 8 at the University of Florida Interdisciplinary Center for Biotechnology Research (UF/ICBR) Cytometry Core Facility (RRID: SCR_019119). Most images were captured with the 20x (0.75 NA, numerical aperture) objective. To image PI-stained seed morphology, fluorescence-lifetime imaging microscopy (FLIM) was conducted using a 63x (1.20 NA) objective with 50 photons/s and post-processing in Leica LAS X. For sequential staining, 0.01% Calcofluor White (CF) was used before PI. Whole-seed immunolabeling was conducted according to (Voiniciuc, 2017), using JIM5 (CarboSource), followed by goat-anti-Rat AlexaFluor488 (Invitrogen), and counter-staining with 0.01% Directed Red 23 (DR23; analogous to Pontamine S4B) in 50 mM NaCl. The following excitation | emission settings were used for CF (405 nm | 450 – 470 nm), green (500 nm | 505 – 550 nm), red (550 nm | 600 – 640 nm) fluorescence.

### Additional seed analyses

Scanning electron microscopy (SEM) of dry seeds was conducted as previously described (Jacobson et al., 2026) at the University of Florida ICBR Electron Microscopy Core Facility (RRID: SCR_019146).

Total mucilage was extracted from seeds using water and a ball mill (30 min at 30 Hz) as previously described (Voiniciuc and Günl, 2016). Matrix polysaccharides were hydrolyzed using 2M trifluoroacetic acid (TFA) and analyzed using High Performance Anion Exchange Chromatography with Pulsed Amperometric Detection (HPAEC-PAD) using an established Metrohm IC Vario system (Jacobson et al., 2026).

### Figure preparation and statistics

Micrographs were processed uniformly post-acquisition in the Fiji distribution of ImageJ (https://imagej.net/software/fiji/). Graphs were prepared in Microsoft Excel, PAST (https://www.nhm.uio.no/english/research/resources/past/), or R package Tidyplots (https://tidyplots.org/). ANOVA analysis was conducted in PAST. Final figures were assembled in Inkscape.

## Results and Discussion

### PEEL1 and PEEL1L are phylogenetically related to RRTs

Phylogenetic analysis of Arabidopsis GT106 proteins (**Fig. 1A**) identified three sub-clades that could affect pectin synthesis: **a)** a pair of uncharacterized genes that we named *PEEL1* (At5g64600) and *PEEL1L* (At1g22460); **b)** the original RRT1-RRT4 isoforms; and c) four additional putative RRTs, including FRB1. Like other GT106 members, PEEL1/1L proteins have a N-terminal transmembrane region followed by a conserved PF10250 domain (**Fig. 1B**). When transiently expressed in *N. benthamiana* leaves, PEEL1-GFP localized in small punctae (**Fig. 1C**) that co-localized with the Golgi marker GmMan1-mCherry (Nelson et al., 2007). In contrast, a GFP-only control showed continuous cytosolic fluorescence. PEEL1-GFP and Golgi marker showed coordinated movement in time-lapse experiments (**Supplementary Video 1**), as expected for the site of matrix polysaccharide biosynthesis.

### *peel1* seeds have reduced mucilage release, despite sufficient pectin

*PEEL1/1L* genes are highly expressed in seeds and other reproductive organs, with transcription peaking in mature pollen (**Supplementary Fig. 1**). Leveraging the GABI-DUPLO collection (Bolle et al., 2013), we obtained single and double mutants (*dm*) for functional analyses (**Fig. 2A**) and confirmed they were homozygous for the reported T-DNA insertions **(Supplementary Fig. 2)**. Each mutation reduced the transcription of the affected gene to trace levels compared to WT (**Fig. 2B**). The *peel1* single mutant and the *dm* seeds showed patchy mucilage release (**Fig. 2C**), unlike the uniform capsules of WT and *peel1L* seeds. The *peel1* RR staining defect was rescued by constitutive *35S:PEEL1* with or without a GFP tag (**Fig. 2C**). Overall, the *peel1* and *dm* mucilage area was ∼40% smaller than WT and *peel1L,* and this reduction was complemented by the *PEEL1* transgene (**Fig. 2D**). In contrast to the mucilage defects, the *peel1/1L* mutants had normal seed area (**Fig. 2E**) and no obvious changes to the development of other plant tissues.

**Figure 2.**
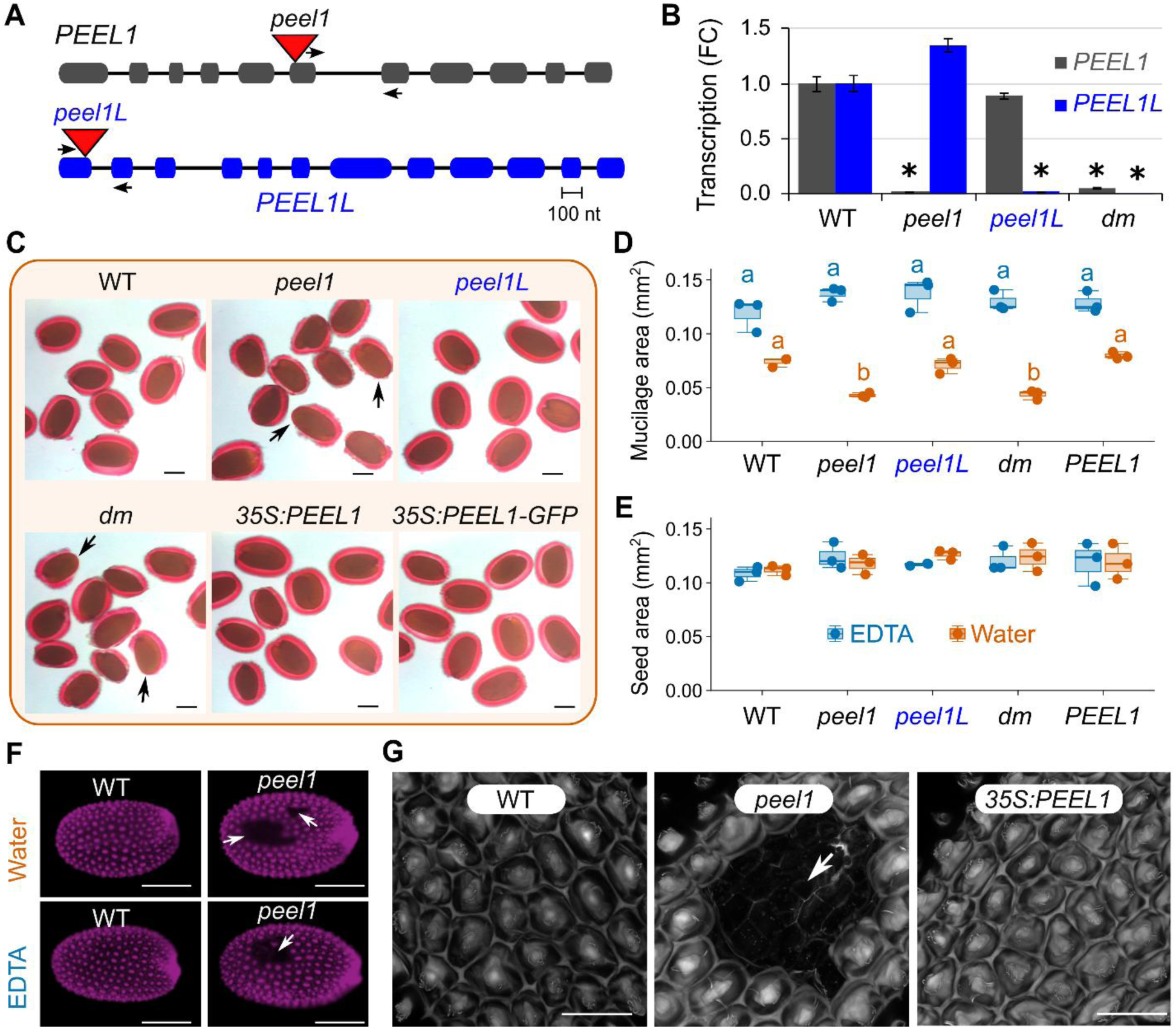
*peel1* alters seed mucilage release. **(A)** Gene structure and characterized mutations. Boxes show exons and lines denote introns. Small arrows indicate qPCR primers. **(B)** Mean + SD (three biological replicates) transcript levels in mature green siliques. Fold change (FC), normalized to housekeeping genes, was set as 1.0 for PEEL1/1L in WT. **(C)** Mucilage RR staining for water-imbibed WT, mutant, and *peel1* complemented seeds. Arrows mark regions with reduced pectin release. **(D)** RR-stained mucilage area and **(E)** seed area using EDTA and water treatments. Box and jitter plots show three biological replicates (∼20 seeds each). Different letters for mucilage area indicate significant differences with one-way ANOVA (Tukey’s post hoc test, *P* < 0.05). **(F)** PI-stained seed epifluorescence, and **(G)** cellular details revealed by PI fluorescence lifetime (FLIM) and Z-stack projections. Arrows show detached or missing epidermal cells. Scale bars = 200 µm **(C, F)** or 50 µm **(G)**. **Alt text:** Figure 2. *peel1* and *peel1L* mutants and their effects on seed mucilage extrusion and epidermal cell adhesion. (A) Gene structures of PEEL1 and PEEL1L, indicating the positions of T-DNA insertions in the respective mutant lines, and the arrows indicate the locations of primers used for gene expression analysis. (B) Bar graph showing the relative expression levels of PEEL1 and PEEL1L in single and double mutant backgrounds. (C) Stereomicroscope images of seed staining illustrating defects in mucilage extrusion in *peel1* single and double mutants; arrows depict the mucilage defects. (D) Jitter and box plot showing the mucilage area in seeds hydrated in water or treated with EDTA and (E) seed area, shown as two different colors for each treatment. (F) Epifluorescence microscope images showing epidermal cell detachment in stained *peel1* single mutants following hydration in water or EDTA treatment. (G) Fluorescence-lifetime imaging microscopy images providing detailed visualization of epidermal cell detachment.

To investigate whether *peel1* has altered pectin gelling properties, dry seeds were pretreated with the 50 mM EDTA to disrupt Ca^2+^ crosslinks prior to staining. The EDTA pretreatment alleviated the patchy mucilage defects of *peel1* and *dm* **(Supplementary Fig. 3)**, resulting in WT-like large capsule areas (**Fig. 2D**). In addition to the *35S*-driven constructs that complemented the *peel1* mucilage defects, *PEEL1* was overexpressed using the stronger *TBA2* promoter, which is seed coat-specific. The *TBA2:PEEL1* transgene (with or without a C-terminal GFP tag) showed a dominant negative effect and impaired mucilage release in *peel1* as well as WT backgrounds (**Supplementary Fig. 4**). Most *TBA2:PEEL1* seeds floated and could not be rescued by EDTA pretreatment, indicating that optimal expression of *PEEL1* is important for mucilage release from seed coat epidermal (SCE) cells.

Since *peel1* disrupts the ability of seed polysaccharides to expand upon hydration, we profiled the composition of mucilage extracts alongside two known pectin mutants. HPAEC-PAD analysis of total mucilage extractable with water did not reveal any changes between WT, *peel1/1L* single mutants, or *dm* seeds in two independent growth batches (**Supplementary Fig. 5**). As previously reported (Voiniciuc et al., 2018), *muci70 and gaut11* produced significantly less Rha and GalA compared to WT, with compensatory changes in the minor components of mucilage. To further investigate the cause for patchy mucilage extrusion from *peel1* and *dm*, PI was used to stain unesterified pectin domains on the seed surface. WT and *peel1L* seeds showed a uniform distribution of PI-stained primary walls, but *peel1* and *dm* seeds had large surface gaps that persisted even after EDTA pretreatment (**Fig. 2F**). Using FLIM confocal microscopy, PI-free regions of *peel1* and *dm* seeds were shown to consist of detached epidermal cell clusters that reveal the underlying primary walls (**Fig. 2G**). The newly exposed layer may correspond to the remainder of the ovule integument layers that are crushed between the epidermis and the endosperm as the seed matures (Haughn and Chaudhury, 2005).

### PEEL1, unlike PEEL1L, maintains cell adhesion in the seed epidermis

Since the *peel1L* mutant resembled the WT, and the *dm* seeds resembled the *peel1* single mutant, we tested if *PEEL1L* overexpression could rescue the *peel1* defects. Unlike equivalent *PEEL1* construct, none of the independent *35S:PEEL1L* transformants altered the *peel1* RR staining defects (**Fig. 3A**). We initially expected the PEEL1/1L paralogs to be functionally redundant, even though their protein sequence identity is lower than other GT106 pairs such as MSR1/2 (**Supplementary Fig. 6**). Upon closer inspection, we discovered that PEEL1L has a polymorphism equivalent to the D390E mutation for FRB1 (**Fig. 3B; Supplementary Fig. 7**). Despite similar PF10250 structures for PEEL1/1L and RRTs in AlphaFold3 (**Supplementary Fig. 8**), the D390E mutation is known to abolish RRT activity *in vitro* and may block GT106 protein binding to nucleotide sugars (Robichaux et al., 2026). Therefore, only PEEL1 is likely to be an active enzyme like RRTs, and we hypothesized that it is specifically contributing to a pectic domain important for primary cell wall adhesion.

**Figure 3.**
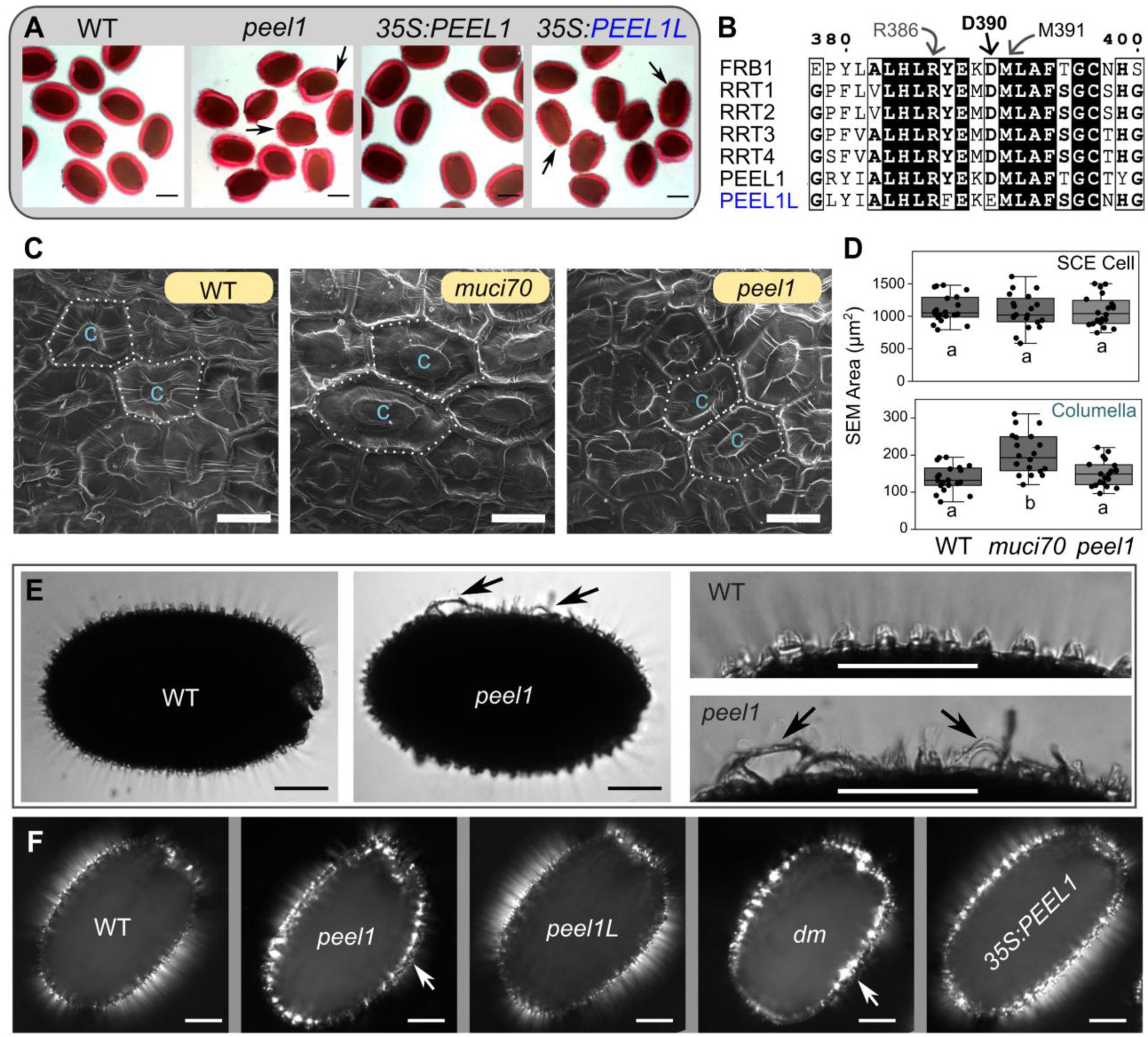
*PEEL1* maintains epidermal wall adhesion in hydrated seeds. **(A)** RR staining showing that *PEEL1L* did not complement *peel1* seeds. **(B)** Alignment of active site region for RRTs and PEEL1/1L. The D390 residue, important for FRB1’s interaction with UDP-Rha, is not conserved in PEEL1L. **(C)** Scanning electron micrographs (SEM) of dry seeds. **(D)** Box and jitter plots of SEM cell and columella areas (20 cells per genotype). Different letters indicate significant differences with one-way ANOVA (Tukey’s post-hoc test, *P* < 0.05). **(E)** Unstained seeds in water with brightfield microscopy. **(F)** Birefringence of crystalline polymers around hydrated seeds. Arrows in **(E)** and **(F)** indicate cell wall adhesions defects. Scale bars = 200 µm **(A)**, 25 µm **(C)**, and 100 µm **(E, F)**. **Alt text:** Figure 3. Characterization of PEEL1L overexpression lines and seed coat cellular defects in PEEL1 mutants. (A) Stereomicroscope images of stained seeds from PEEL1L overexpression lines, illustrating mucilage extrusion phenotypes. (B) Partial protein sequence alignment of PEEL1/1L and characterized RRT proteins, highlighting a mutation affecting a conserved amino acid in PEEL1L. (C) SEM images of seed coat epidermal cells in WT, *muci70*, and *peel1* mutant seeds. (D) Jitter and box plots quantifying seed coat epidermal cell and columella areas in WT and mutant seeds. (E) Transmitted-light microscopy images of hydrated, unstained *peel1* mutant seeds compared with WT, with arrows indicating epidermal cell detachment. (F) Micrographs with arrows showing defects in birefringence in *peel1* single and double mutants.

To test this hypothesis, the dry seed surface morphology was assessed with SEM. The *peel1* SCE cells resembled the WT, while *muci70* seeds had larger columellae due to reduced mucilage accumulation (**Fig. 3C, D**). The *peel1* SCE cells showed primary wall adhesion defects post-hydration with brightfield microscopy (**Fig. 3E**) or under polarized light (**Fig. 3F**). Even without staining, regions of crystalline wall fibers were observed in clusters close to the surface of *peel1* and *dm* seeds, with “suspended bridges” forming between some SCE cells.

### *peel1* disrupts (hemi)cellulosic fibers and pectin distribution

Mucilage architecture was further assessed after dual staining with PI and CF, which binds. Unlike the uniform CF-stained rays around WT and *peel1L* seeds, *peel1* and *dm* mucilage had clusters of missing or mislocalized β-glucans (**Fig. 4A**). The disruption of CF-stained rays was most severe, but not restricted, to *peel1* or *dm* regions lacking PI staining. These observations were consistent with the overall changes in birefringence (**Fig. 3F**). Furthermore, pectin was immunolabelled with the JIM5 antibody and crystalline cellulose was counter-stained with DR23. JIM5 binds to low-methylesterified HG close to the seed surface (Griffiths et al., 2014), and uniformly labelled SCE wall outlines for WT and *peel1L* (**Fig. 4B**). In contrast, most *peel1* and *dm* SCE cells had severe perturbations in JIM5 epitopes, with larger pieces of detached walls instead of thin rays (**Fig. 5**). As observed for RR staining (**Fig. 2C-E**), the *35S*:*PEEL1* transgene fully restored glucan and HG distribution around the seed surface (**Fig. 4A,B**).

**Figure 4.**
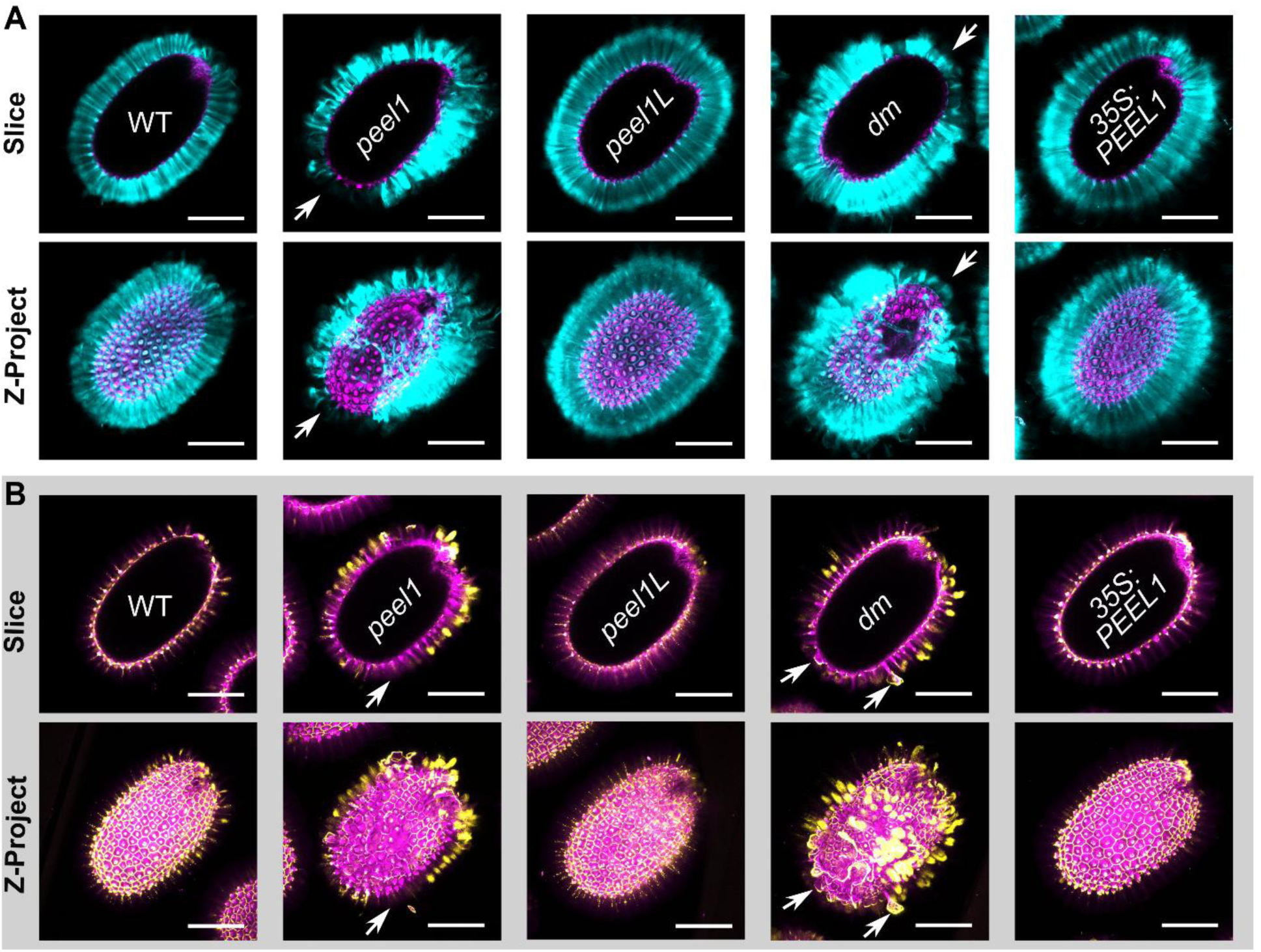
Loss of PEEL1 disrupts β-glucan and HG polymer distribution. **(A)** Single optical sections and Z-projections of β-glucan labelled with calcofluor (cyan). PI (magenta) was used as a counterstain for unesterified pectin. **(B)** Immunolabeling of low-methylesterified HG using the JIM5 antibody (yellow). Optical sections and Z-projections show JIM5 signal overlayed with DR23 counterstaining of crystalline cellulose (magenta). White arrows in **(A)** and **(B)** indicate regions with missing or severely disrupted polysaccharides. Scale bars = 200 µm. **Alt text:** Figure 4. Confocal microscopy analysis of polysaccharide organization and low-methylesterified pectin distribution in seed mucilage. (A) Confocal microscopy images of seeds co-stained with propidium iodide (magenta) and calcofluor white (cyan), shown as single optical sections and Z-projections. (B) Confocal microscopy images of seeds immunolabeled to detect low-methylesterified pectin (yellow) and counterstained with DR23 (magenta), shown as single optical sections and Z-projections. White arrows indicate regions of altered polysaccharide organization or disruption within the seed mucilage.

## Conclusions

We propose that Golgi-localized PEEL1 is required for the synthesis of a pectin domain facilitates adhesion within and/or between SCE primary walls. RG-I is the most likely candidate due to i) its importance for cell-cell adhesion in plants, including woody tissues (Yang et al., 2020), and ii) the phylogenetic proximity of PEEL1 to active RRTs (**Fig. 1A**, Takenaka et al., 2018; Robichaux et al., 2026). Despite no changes in overall mucilage composition (**Supplemental Fig. 5**), primary wall components were disorganized around *peel1* seed surfaces. Bright patches of low-methylesterified HG for *peel1* seeds are consistent with increased pectin-Ca^2+^ crosslinking (**Fig. 4B**) and the ability of a cation chelator to restore mucilage expansion (**Fig. 2D, Supplemental Fig. 3**). Even after EDTA pre-treatment (**Fig. 2F,G**), *peel1* still had regions with detached SCE cells, reminiscent of the cellular cracks reported for the *ruby* galactose oxidase mutant (Šola et al., 2019). Nevertheless, the *peel1* mutant presents a unique phenotype among known mucilage mutants (Voiniciuc et al., 2015c), with *ruby* showing more severe staining defects and significant changes in the arabinogalactan branching on RG-I (Šola et al., 2019). Other known mutants that increase pectin gelation (*sbt1.7*, Rautengarten et al., 2008; *per36*, Kunieda et al., 2013; pmei6, Saez-Aguayo et al., 2013; *fly1*, Voiniciuc et al., 2013) further limit mucilage release upon hydration and show detached primary cell walls as large sheets or numerous individual discs (only *fly1*). In contrast, the distribution of pectin in primary walls, rather than its abundance in mucilage, is clearly patchy when the *PEEL1* expression is knocked out (**Fig. 2-4**). Altered deposition of pectin in *peel1* could then impair the organization of β-glucans, and interfere with the rupture of primary walls important for mucilage release. Unlike its closest paralog, *PEEL1L* did not affect seed coat phenotypes and cannot encode an active RRT due to the loss of the conserved D390 residue. Although future studies are required to determine whether PEEL1 uses UDP-Rha like FRB1 or acts on other carbohydrates, our findings indicate that different GT106 family members (**Fig. 1A**) contribute to cell-cell adhesion in distinct tissues. Presence of numerous RRT-like proteins in a single species may reflect a requirement for specialized enzymes that generate structurally distinct pectin domains tailored to the adhesive needs of specific wall types.

## Author contributions

A.G.V. led most of the experiments, data acquisition, data analyses, and manuscript drafting. T.J. led the confocal microscopy experiments. C.V. designed the project, provided supervision, analyzed data and revised the manuscript.

## Acknowledgements

We thank Bo Yang for initial mutant seed propagation, Lianna Larson for technical assistance with staining and carbohydrate analysis at the start of the project, and Karen Kelley for the preparation of SEM samples.

## Supplementary Material

Supplementary material is available online.

**Supplementary Table 1.** Sequences of primers used for cloning, genotyping, and qPCR.

**Supplementary Table 2.** Modular cloning parts for plasmid assembly.

**Supplementary Figure 1.** *PEEL1* and *PEEL1L* transcriptional profiles.

**Supplementary Figure 2**. Genotyping of mutations in PEEL1/1L.

**Supplementary Figure 3.** Effects of a cation chelator on seed mucilage release.

**Supplementary Figure 4.** Promoter strength controls *PEEL1*-dependent mucilage release.

**Supplementary Figure 5.** Mutations in *PEEL1/1L* do not alter total mucilage composition.

**Supplementary Figure 6.** Percent identity matrix for selected Arabidopsis GT106 proteins.

**Supplementary Figure 7.** Multiple sequence alignment for RRTs and PEEL1/1L.

**Supplementary Figure 8.** AlphaFold3 models of GT domain for PEEL1/1L.

**Supplementary Video 1**. Timelapse of PEEL1 co-localization with a Golgi marker.

## Data availability statement

Data is provided in the main manuscript and supplementary section. Additional information is available upon request from the corresponding author.

## References

Amos RA, Atmodjo MA, Huang C, Gao Z, Venkat A, Taujale R, Kannan N, Moremen KW, Mohnen D (2022) Polymerization of the backbone of the pectic polysaccharide rhamnogalacturonan I. Nat Plants 8: 1289–1303

Bolle C, Huep G, Kleinbölting N, Haberer G, Mayer K, Leister D, Weisshaar B (2013) GABI - DUPLO: a collection of double mutants to overcome genetic redundancy in *A rabidopsis thaliana*. The Plant Journal 75: 157–171

Cosgrove DJ (2024) Structure and growth of plant cell walls. Nat Rev Mol Cell Biol 25: 340– 358

Engler C, Youles M, Gruetzner R, Ehnert T-M, Werner S, Jones JDG, Patron NJ, Marillonnet S (2014) A Golden Gate Modular Cloning Toolbox for Plants. ACS Synth Biol 3: 839–843

Griffiths JS, Tsai AYL, Xue H, Voiniciuc C, Šola K, Seifert GJ, Mansfield SD, Haughn GW (2014) SALT-OVERLY SENSITIVE5 mediates arabidopsis seed coat mucilage adherence and organization through pectins. Plant Physiology 165: 991–1004

Haughn G, Chaudhury A (2005) Genetic analysis of seed coat development in Arabidopsis. Trends in Plant Science 10: 472–477

Jacobson T, Edwards M, Qiande M, Robert M, Moncrieff J, Voiniciuc C (2026) Golgi-localized mannanases sustain hemicellulose biosynthesis. New Phytologist 250: 834–844

Kunieda T, Shimada T, Kondo M, Nishimura M, Nishitani K, Hara-Nishimura I (2013) Spatiotemporal Secretion of PEROXIDASE36 Is Required for Seed Coat Mucilage Extrusion in Arabidopsis. Plant Cell 25: 1355–1367

McGee R, Dean GH, Mansfield SD, Haughn GW (2019) Assessing the utility of seed coat-specific promoters to engineer cell wall polysaccharide composition of mucilage. Plant Mol Biol 101: 373–387

Nelson BK, Cai X, Nebenführ A (2007) A multicolored set of *in vivo* organelle markers for co-localization studies in Arabidopsis and other plants. The Plant Journal 51: 1126–1136

Neumetzler L, Humphrey T, Lumba S, Snyder S, Yeats TH, Usadel B, Vasilevski A, Patel J, Rose JKC, Persson S, et al (2012) The FRIABLE1 Gene Product Affects Cell Adhesion in Arabidopsis. PLoS ONE 7: e42914

Rautengarten C, Usadel B, Neumetzler L, Hartmann J, Büssis D, Altmann T (2008) A subtilisin-like serine protease essential for mucilage release from Arabidopsis seed coats. Plant Journal 54: 466–480

Robichaux KJ, Blea MN, Amos RA, Huang C, Mohnen D, Wallace IS (2026) A characterization of recombinant Arabidopsis FRIABLE1 (FRB1) reveals robust rhamnogalacturonan-I rhamnosyltransferase activity and critical catalytic residues. Journal of Biological Chemistry 302: 111305

Saez-Aguayo S, Ralet M-C, Berger A, Botran L, Ropartz D, Marion-Poll A, North HM (2013) PECTIN METHYLESTERASE INHIBITOR6 Promotes Arabidopsis Mucilage Release by Limiting Methylesterification of Homogalacturonan in Seed Coat Epidermal Cells. The Plant Cell 25: 308–323

Saez-Aguayo S, Sanhueza D, Jara V, Galleguillos B, De La Rubia AG, Largo-Gosens A, Moreno A (2025) Mucilicious methods: Navigating the tools developed to Arabidopsis Seed Coat Mucilage analysis. The Cell Surface 13: 100134

Šola K, Gilchrist EJ, Ropartz D, Wang L, Feussner I, Mansfield SD, Ralet M-C, Haughn GW (2019) RUBY, a Putative Galactose Oxidase, Influences Pectin Properties and Promotes Cell-To-Cell Adhesion in the Seed Coat Epidermis of Arabidopsis. Plant Cell 31: 809–831

Takenaka Y, Kato K, Ogawa-Ohnishi M, Tsuruhama K, Kajiura H, Yagyu K, Takeda A, Takeda Y, Kunieda T, Hara-Nishimura I, et al (2018) Pectin RG-I rhamnosyltransferases represent a novel plant-specific glycosyltransferase family. Nature Plants 4: 669–676

Voiniciuc C (2017) Whole-seed Immunolabeling of Arabidopsis Mucilage Polysaccharides. BIO-PROTOCOL. doi: 10.21769/BioProtoc.2323

Voiniciuc C, Dama M, Gawenda N, Stritt F, Pauly M (2019) Mechanistic insights from plant heteromannan synthesis in yeast. Proc Natl Acad Sci USA 116: 522–527

Voiniciuc C, Dean GH, Griffiths JS, Kirchsteiger K, Hwang YT, Gillett A, Dow G, Western TL, Estelle M, Haughn GW (2013) FLYING SAUCER1 Is a Transmembrane RING E3 Ubiquitin Ligase That Regulates the Degree of Pectin Methylesterification in Arabidopsis Seed Mucilage. The Plant Cell 25: 944–959

Voiniciuc C, Engle KA, Günl M, Dieluweit S, Schmidt MH-W, Yang J-Y, Moremen KW, Mohnen D, Usadel B (2018) Identification of Key Enzymes for Pectin Synthesis in Seed Mucilage. Plant Physiol 178: 1045–1064

Voiniciuc C, Guenl M, Schmidt MH-W, Usadel B (2015a) Highly Branched Xylan Made by IRX14 and MUCI21 Links Mucilage to Arabidopsis Seeds. Plant Physiol pp.01441.2015

Voiniciuc C, Günl M (2016) Analysis of Monosaccharides in Total Mucilage Extractable from Arabidopsis Seeds. BIO-PROTOCOL 6: 1–12

Voiniciuc C, Schmidt MH-W, Berger A, Yang B, Ebert B, Scheller HV, North HM, Usadel B, Günl M (2015b) MUCILAGE-RELATED10 Produces Galactoglucomannan That Maintains Pectin and Cellulose Architecture in Arabidopsis Seed Mucilage. Plant Physiol 169: 403– 420

Voiniciuc C, Yang B, Schmidt M, Günl M, Usadel B (2015c) Starting to Gel: How Arabidopsis Seed Coat Epidermal Cells Produce Specialized Secondary Cell Walls. IJMS 16: 3452–3473

Wang Y, Mortimer JC, Davis J, Dupree P, Keegstra K (2013) Identification of an additional protein involved in mannan biosynthesis. Plant Journal 73: 105–117

Weber E, Engler C, Gruetzner R, Werner S, Marillonnet S (2011) A Modular Cloning System for Standardized Assembly of Multigene Constructs. PLoS ONE 6: e16765

Yang H, Benatti MR, Karve RA, Fox A, Meilan R, Carpita NC, McCann MC (2020) Rhamnogalacturonan-I is a determinant of cell–cell adhesion in poplar wood. Plant Biotechnology Journal 18: 1027–1040

Zhong R, Phillips DR, Adams ER, Ye Z-H (2023) An Arabidopsis family GT106 glycosyltransferase is essential for xylan biosynthesis and secondary wall deposition. Planta 257: 43

