## Supplemental Tables and Figures for "PEELING WALLS1 encodes a GT106 protein required for seed surface integrity"

**Supplementary Table 1.** Sequences of primers used for cloning, genotyping, and qPCR.

| TARGET | FORWARD (5' – 3') | REVERSE (5' – 3') |
|---|---|---|
| GABI-KAT T-DNA (o8474) | – | ataataacgctgcggacatctacatttt |
| *peel1-1* | tcattcctgagctggataagc | acaaaggccttcctctctctg |
| *peel1L-1* | tcacgtgacccttctctaacc | cggtgttgttgatgtgttcc |
| *PEEL1 qPCR* | gcaattgaaagccttggacagaag | agtgagaccataggtgcaaccag |
| *PEEL1L qPCR* | actctgttttctcaccgtacac | cttgggattggagatgagagg |
| *GAPC1 qPCR* | tcagactcgagaaagctgctac | gatcaagtcgaccacacgg |
| *UBQ5 qPCR* | gacgcttcatctcgtcc | ccacaggttgcgttag |
| MoClo backbone | gagtcagtgagcgaggaagc | cctttcgtccactgaagagc |
| *AtPEEL1* optimized cds genotyping | aagccaacatgtccttctcg | – |
| *35S* genotyping | gaggagcatcgtggaaaaag | |
| *TBA2* genotyping | gatcttgggcaagagaggtg | – |
| *tOCS* genotyping | – | ttaggtttgaccggttctgc |
| GFP genotyping | – | aagtcgtgctgcttcatgtg |
| *AtPEEL1* with STOP codon | actgaagacgcAATGgtgaagcacagaaactcatctcg | gatgaagacgtAAGCgcttagggttctttgtgctcctt |
| *AtPEEL1L* with STOP codon | atagaagacatAATGcggcggcggagaaagacggtt | gatgaagacgtAAGCgcttatctcgctttacttggatctctctg |

**Supplementary Table 2.** Modular cloning parts for plasmid assembly. The second part of the table shows the optimized *AtPEEL1* coding sequence synthesized for complementation analyses.

| Source | DNA part | Plasmid ID |
|---|---|---|
| Addgene MoClo Kit 1000000044 | L0 backbone | pICH41308 |
| Addgene MoClo Kit 1000000044 | L0 backbone for C-terminal tagging | pAGM1287 |
| Addgene MoClo Kit 1000000044 | L1 backbone | pICH47742 |
| 1000000047 | p35s short | pICH51277 |
| 1000000047 | Green fluorescent protein | pICSL50008 |
| Addgene MoClo Kit 1000000044 | LM backbone | pAGM8031 |
| Addgene MoClo Kit 1000000044 | M Linker (end-link 2) | pICH50881 |
| **Synthesized *PEEL1* codon-optimized sequence** | | |
| ATGGTGAAGCACAGAAACTCATCTCGTTCGATAATCTCATACTCTTCTTCCATTGCAAGA<br>TTCTTCTCTAGAAAAGCTATTTCTCTCTACTTGATTTTCGTCTTTGCTTTTACCATCTGG<br>GTTCTGGTCTTTAGCTCCAGAAACATTCAAACCGATGATGACCACACCAAACATCAACAA<br>CAACATCATCGGGACCTAATCGACTCCGAATCCTTTCCGCCACCGTATTTGCCTCCTAGG<br>AAGAATTTGCAGAAACCATATGAAAATACTCAACTTTGGACTCCTCCTTTCAGCTTTGGG<br>TTGCATCCATGTGTAAAACCTACTCCCAAGTACAAAGAATTTTCAGAATCAGATCACTAT<br>ATAACAGTGAAAAGTAACGGTGGATTGAATCAGATGCGTACTGGTATCGCAGATATAGTT<br>GCTGTTGCGCACATCATGAACGCCACCTTAGTCATTCCTGAGTTGGATAAGCGATCGTTT<br>TGGCAAGATTCAAGTGTTTTTTCCGATATTTTTGACGAGGAACAATTCATTAAATCTTTG<br>CGAAGAGATGTCAAGGTTATTAAAAAGTTGCCAAAGGAAGTGGAATCGCTACCCAGAGCC<br>AGGAAGCATTTCACGTCTTGGTCTAGTGTTGGCTACTATGAGGAAATGACACACTTGTGG<br>AAGGAGTACAAGGTCATCCATGTCGCAAAATCAGATTCTAGACTTGCTAACAATGACCTG<br>CCTATCGACGTTCAAAGACTGAGATGTCGTGTACTATACCGTGGTTTATGCTTCTCTCCT<br>GCCATTGAAAGCCTTGGACAGAAGTTAGTTGAGAGACTCAAGTCAAGAGCTGGTCGTTAT<br>ATTGCCTTGCACCTGAGATACGAGAAGGACATGTTGGCTTTCACTGGTTGCACCTATGGT<br>CTGACCGACGCTGAATCCGAAGAACTGAGAGTAATGCGGGAAAGTACAAGCCATTGGAAG<br>ATTAAGAGTATAAATTCAACAGAGCAGAGAGAGGAAGGCCTTTGTCCATTGACTCCAAAA<br>GAAGTAGGAATATTTCTGAAAGGTCTAGGATATTCTCAGTCCACAGTCATATACATTGCC<br>GCAGGGGAGATCTACGGCGGTGATGATAGACTCTCTGAGCTTAAGTCGCGCTTCCCAAAC<br>TTAGTTTTCAAGGAAACGCTTGCTGGTAACGAGGAGTTAAAAGGTTTCACTGGCCATGCT<br>ACTAAGACGGCTGCTTTAGATTACATTATTTCTGTTGAGAGTGATGTGTTTGTTCCTTCC<br>CATTCTGGAAACATGGCCAGAGCAGTTGAAGGTCACCGAAGATTTCTTGGACATCGCAGG<br>ACTATCACACCCGACAGGAAAGGATTAGTGAAATTGTTCGTTAAGATGGAGAGAGGTCAG<br>CTAAAAGAAGGACCAAAGTTGTCCAATTTTGTGAATCAAATGCATAAAGACAGACAAGGT<br>GCACCTAGGCGAAGGAAAGGACCAACCCAGGGGATCAAAGGACGTGCTCGTTTTAGAACT<br>GAAGAAGCCTTTTATGAGAATCCATATCCAGAGTGTATTTGCAGTTCAAAGGAGCACAAA<br>GAACCC | | |

**Supplementary Figure 1.** *PEEL1* and *PEEL1L* transcriptional profiles. Expression levels during plant **(A)** and seed **(B)** development. Expression levels are based on the Arabidopsis eFP browser data.

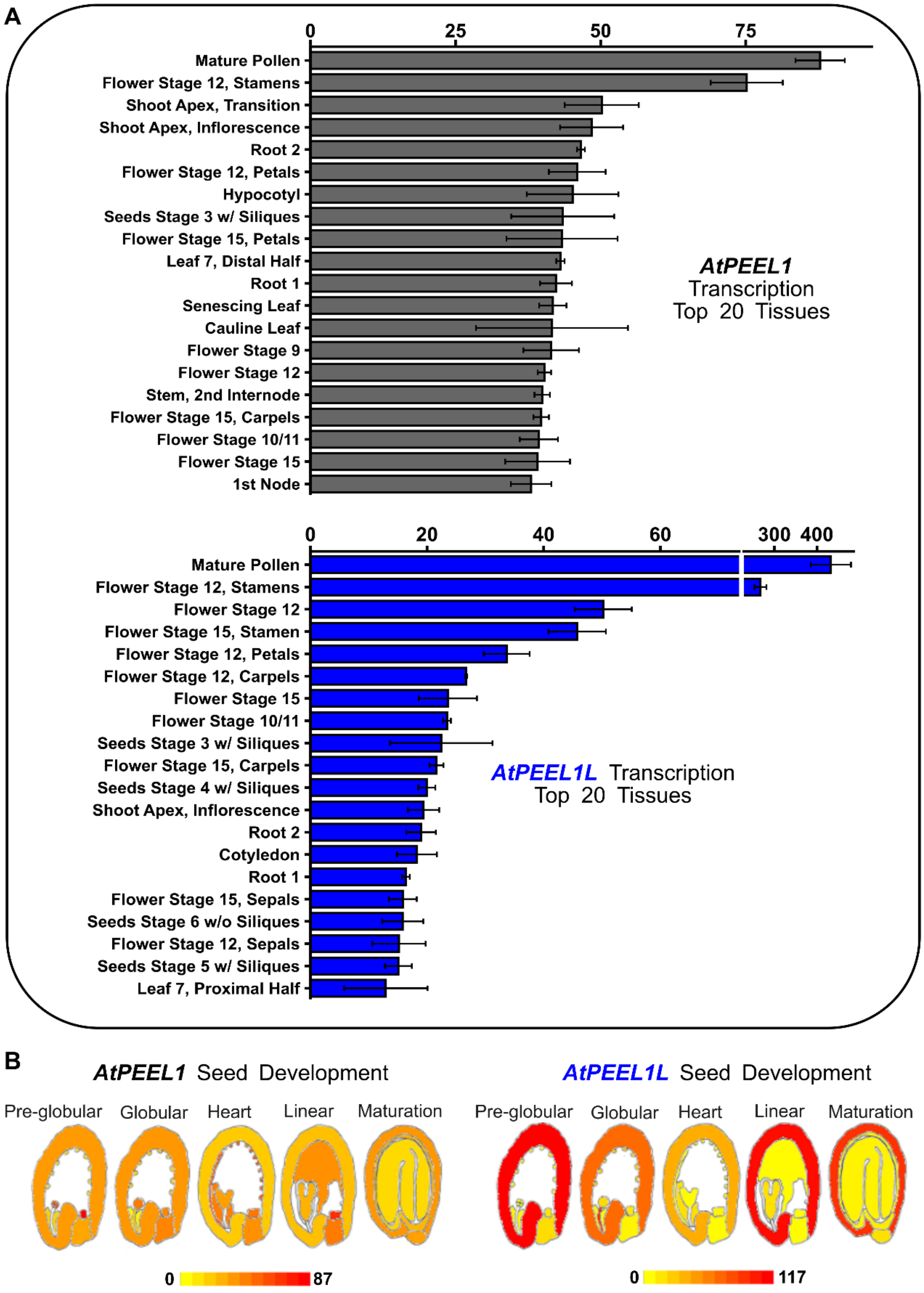

**Supplementary Figure 2**. Genotyping of mutations in PEEL1/1L. **(A)** Gene maps showing the position of T-DNA insertions and of T-DNA genotyping primers (black arrows). **(B)** Genotyping of *peel1,* **(C)** *peel1L,* and **(D)** double mutant (*dm*) plants. PCR reactions were performed by using gene-specific F+R and mutant-specific F+o8474 combinations. Relevant fragment sizes are noted relative to the VWR 100 bp DNA ladder.

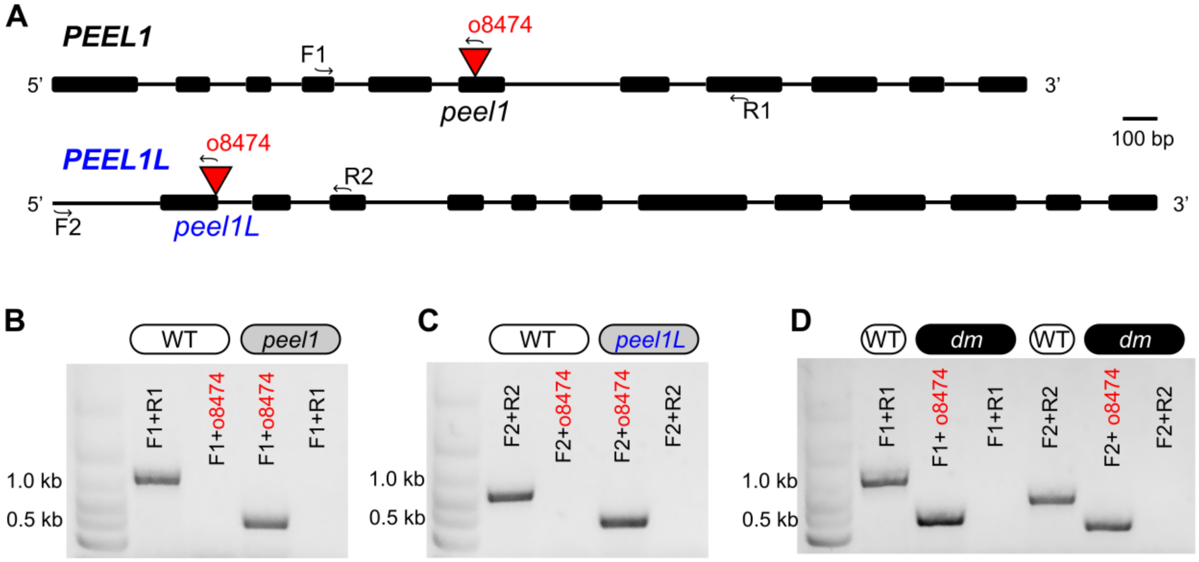

**Supplementary Figure 3**. Effects of a cation chelator on seed mucilage release. RR staining in WT and mutant seeds after initial hydration in water **(A)** or EDTA **(B)**. Images show representative examples of seeds used for the quantification of mucilage and seed areas **(Fig. 2D,E)**. Scale bars = 200 µm.

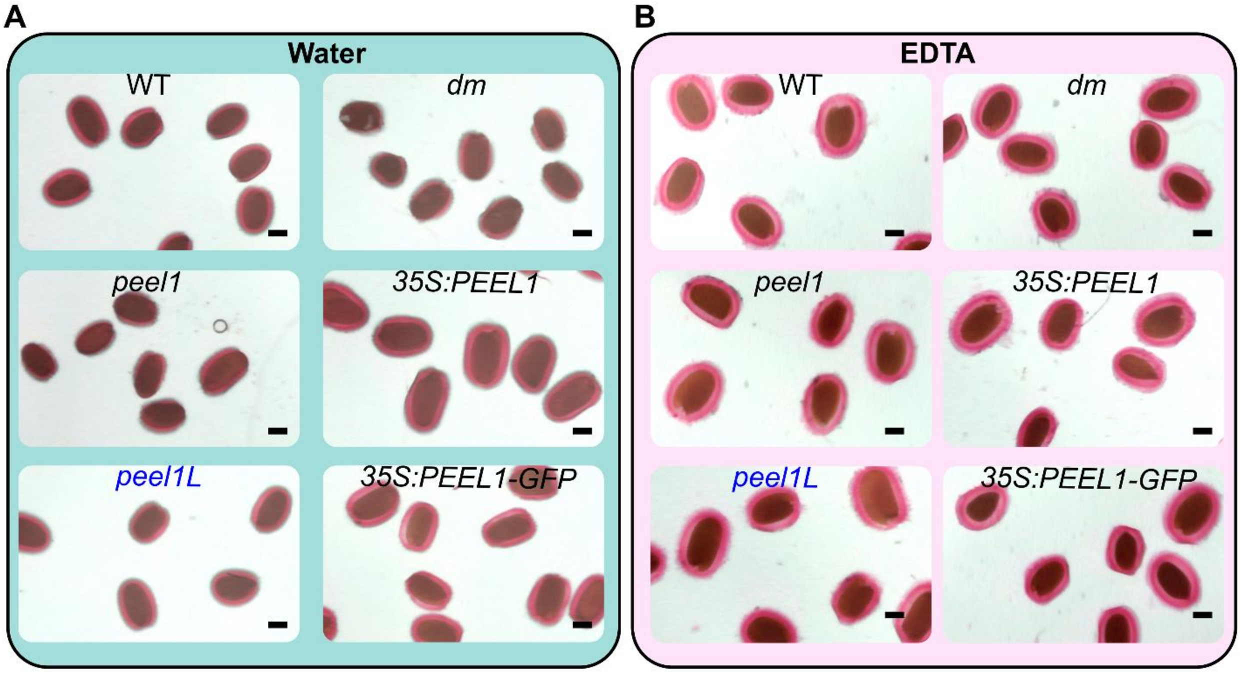

**Supplementary Figure 4**. Promoter strength controls *PEEL1*-dependent mucilage release. **(A)** Seed phenotype of *PEEL1* or *PEEL1-GFP* expressed under the constitutive *35S* or the stronger *TBA2* promoter. *TBA2*:*PEEL1* constructs. Black arrows show that *TBA2:PEEL1* (± C-terminal GFP tag) reduced mucilage release in both WT and *peel1* backgrounds. **(B)** Pre-treatment of seeds with EDTA partially restored mucilage release from *TBA2:PEEL1* overexpression seeds (red arrows). Scale bars = 200 μm.

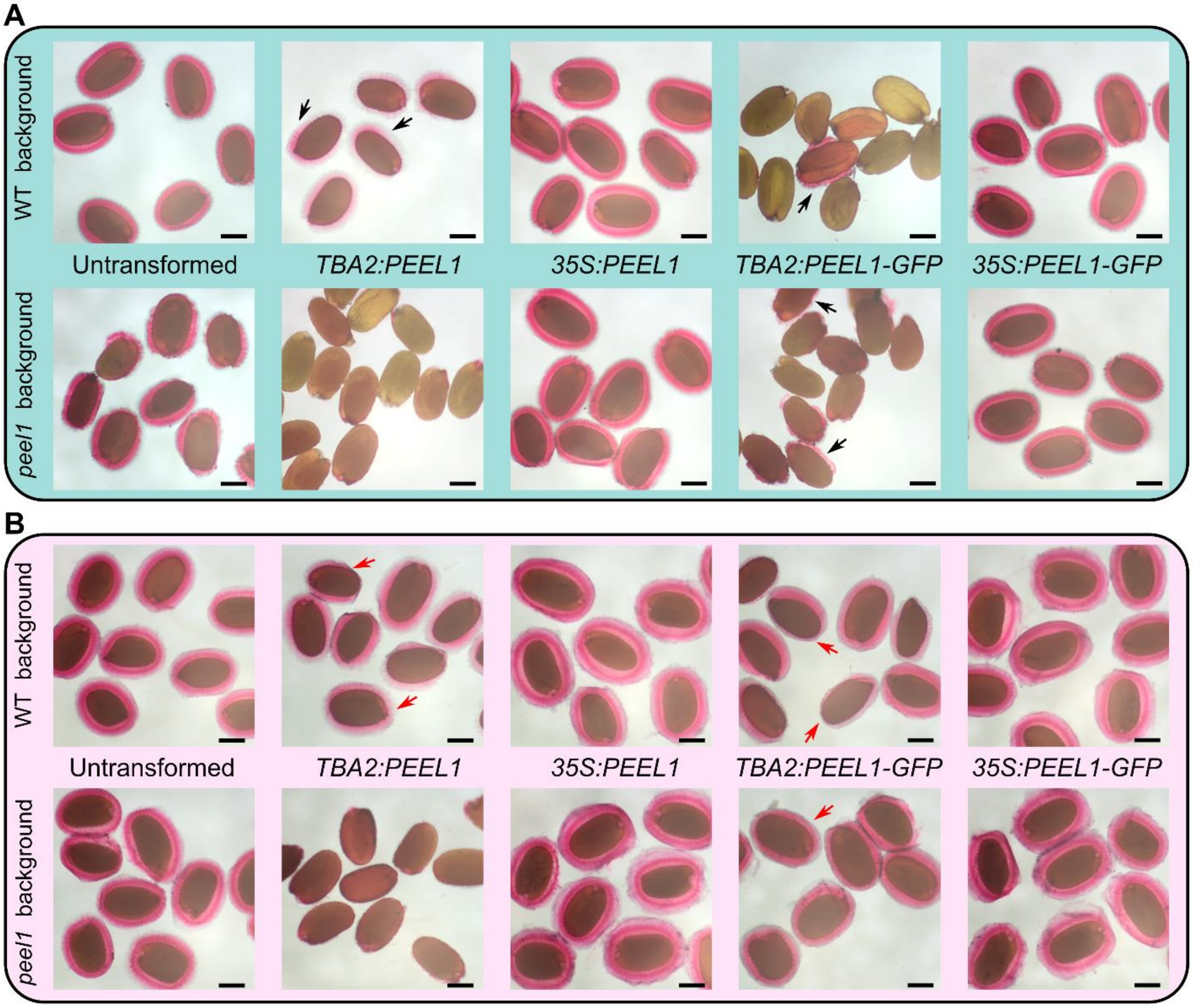

**Supplementary Figure 5.** Mutations in *PEEL1/1L* do not alter total mucilage composition. **(A)** and **(B)** Total seed mucilage extractable with water in two independent growth batches. Bars represent the mean values of four biological replicates (shown as jitters). Arrows show significant increase or decrease compared to WT (two-sided t-test, $P < 0.001$, assuming equal variance). The *muci70* and *gaut11* mutants are included as controls known to have reduced pectin production.

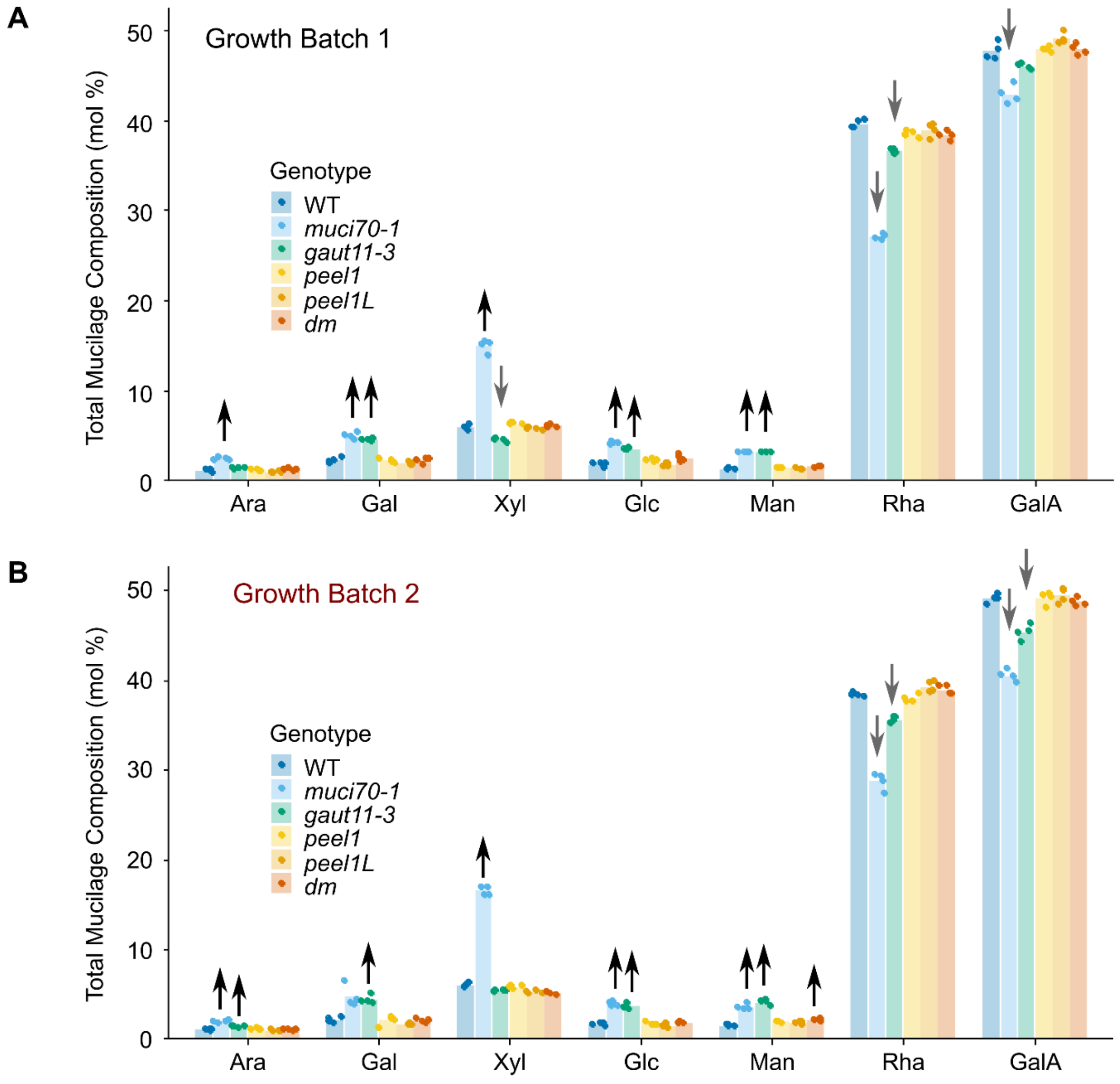

**Supplementary Figure 6.** Percent identity matrix for selected Arabidopsis GT106 proteins. The focus is on RRT-like proteins, with MSR proteins as more distant controls. Values were obtained using multiple sequence alignment in Clustal Omega (https://www.ebi.ac.uk/jdispatcher/msa/clustalo).

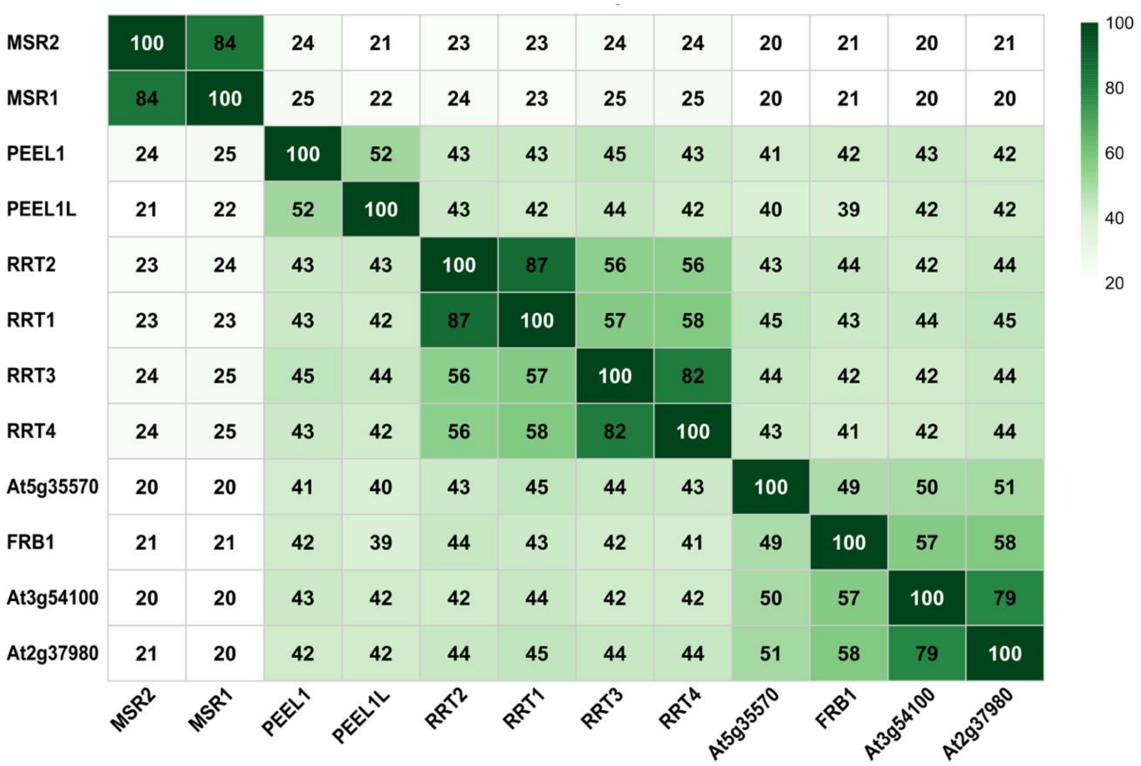

**Supplementary Figure 7.** Multiple sequence alignment for RRTs and PEEL1/1L. The figure was prepared using ESPript and the Muscle algorithm. Regions shaded in black are fully conserved, while black outlines mark partly conserved residues. Amino acids are numbered based on FRB1, and red outlines denote positions important for RRT activity. The […] indicates alignment regions that are not shown to save space.

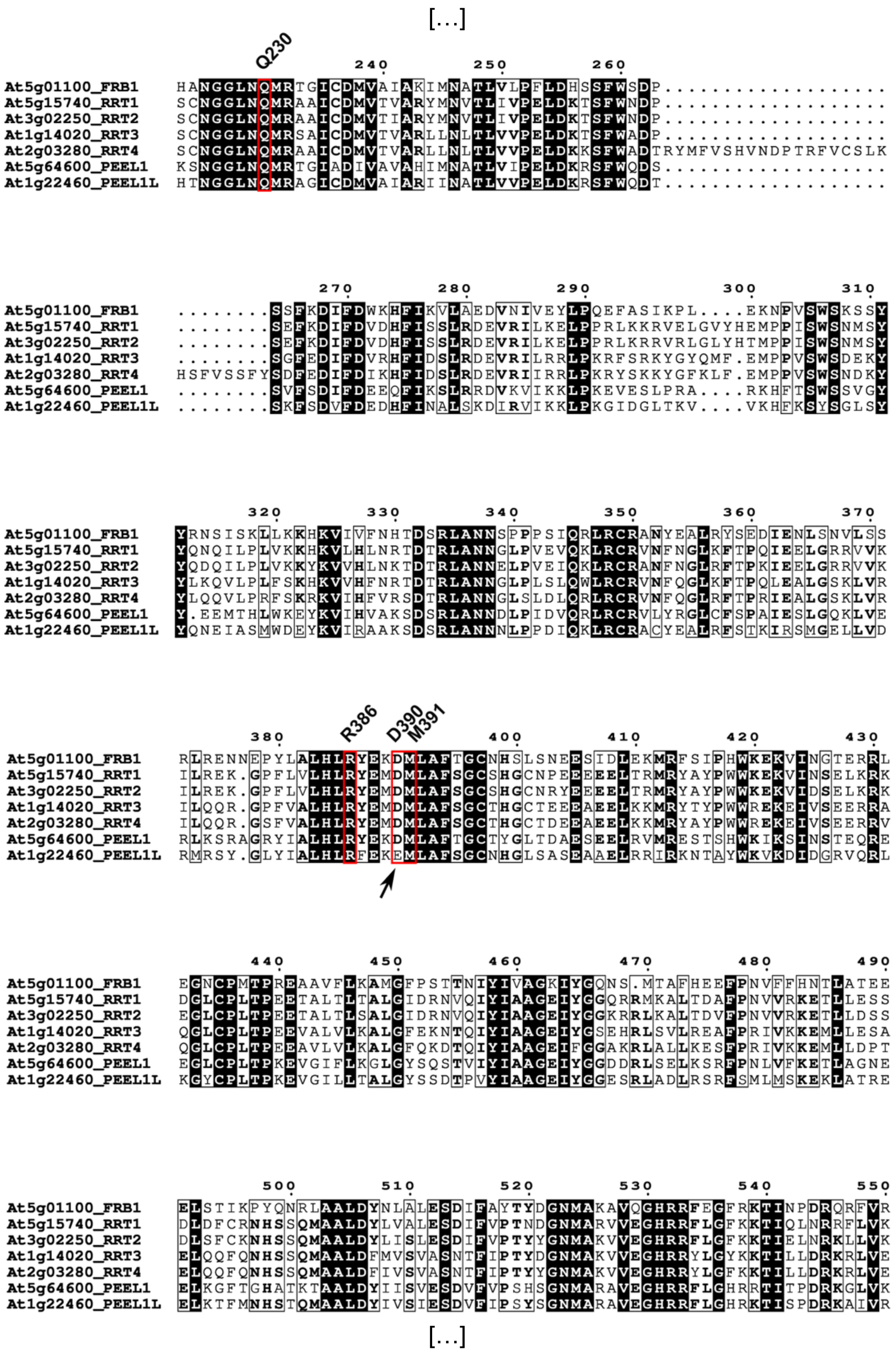

**Supplementary Figure 8.** AlphaFold3 models of GT domain for PEEL1/1L. The folding of the PF10250 domain of **(A)** FRB1 and PEEL1, **(B)** PEEL1 and PEEL1L, and **(C)** FRB1 and PEEL1. Magnified view in **(C)** shows the effects of the D390E change between FRB1 and PEEL1L.

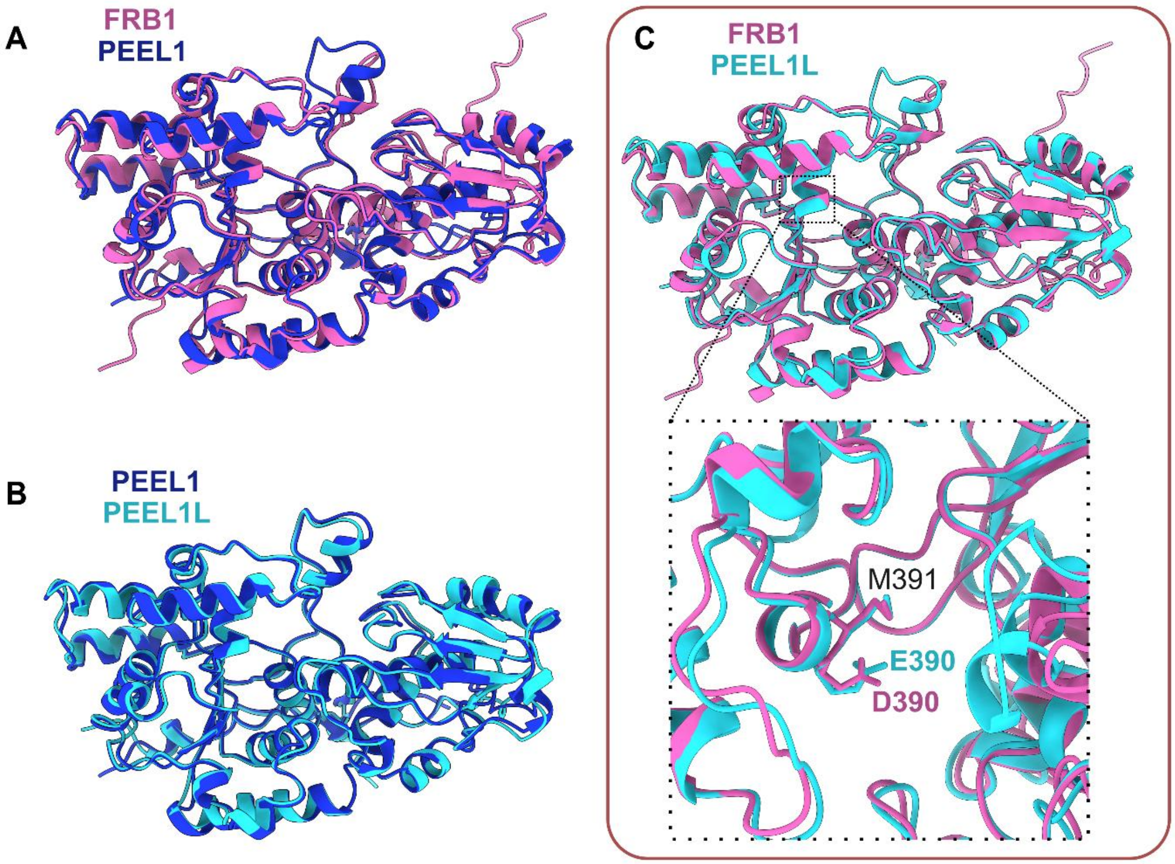
