## Supplemental Video 1 for "PEELING WALLS1 encodes a GT106 protein required for seed surface integrity"

### Slide 1
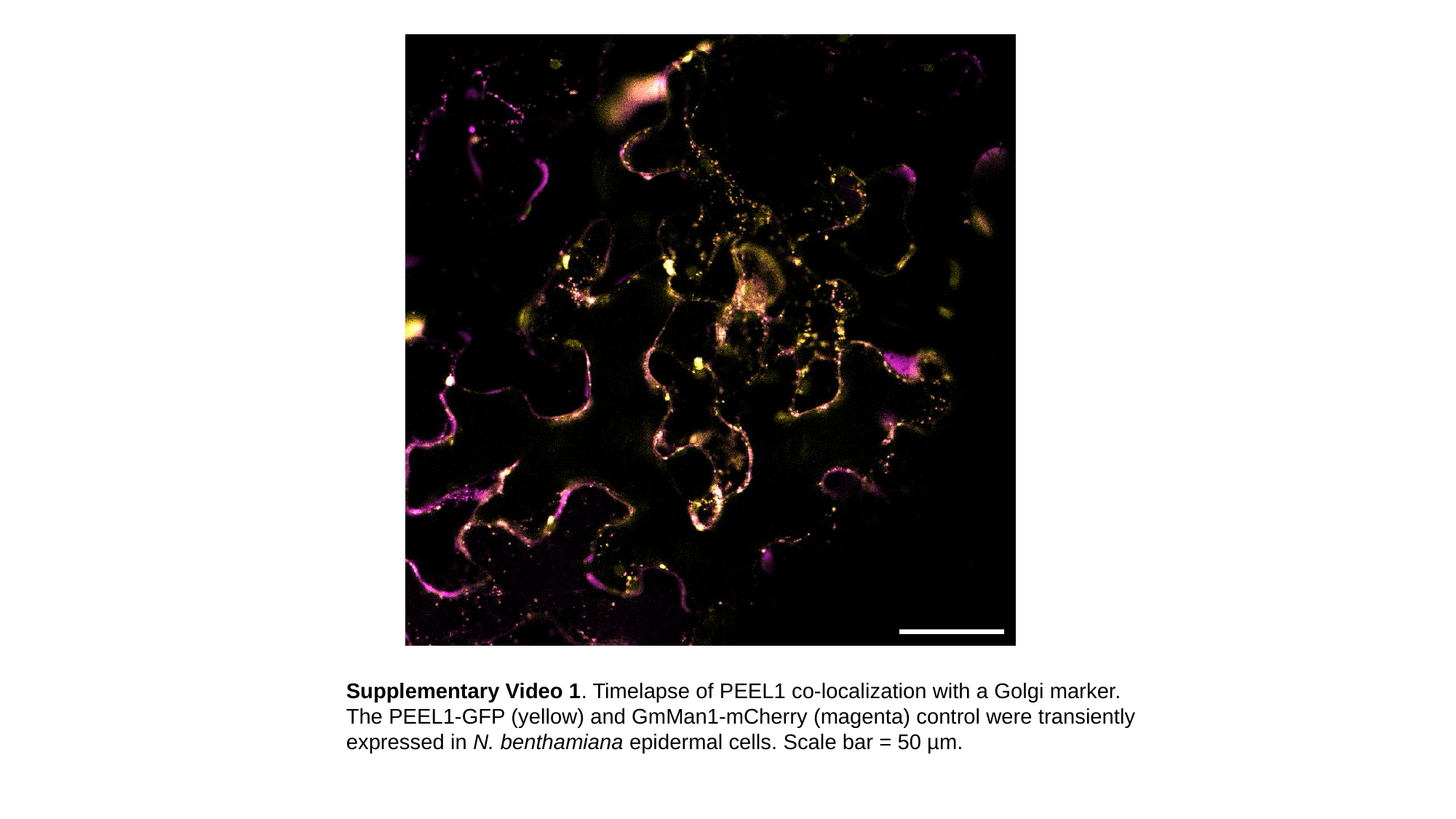

Supplementary Video 1. Timelapse of PEEL1 co-localization with a Golgi marker. The PEEL1-GFP (yellow) and GmMan1-mCherry (magenta) control were transiently expressed in N. benthamiana epidermal cells. Scale bar = 50 µm.
